# Evaluate the efficiency of Mr BMT in macrophage replacement of peripheral organs

**DOI:** 10.1101/2024.06.22.600056

**Authors:** Jinjin Du, Baozhi Yang, Yang He, Yanxia Rao

## Abstract

The successful replacement of endogenous tissue-resident macrophages with healthy bone marrow-derived cells through transplantation presents a promising therapeutic approach for treating disease associated with macrophage dysfunction. Our previous development of the novel microglia replacement by bone marrow transplantation (Mr BMT) method achieved high engraftment efficiency in the central nervous system (CNS), overcoming the limitation of traditional bone marrow transplantation (tBMT). In this study, we aimed to investigate the potential impacts of Mr BMT approach on peripheral organs. The Mr BMT approach involves depleting resident microglia through colony stimulating factor-1 receptor (CSF-1R) inhibition, so we first evaluated the survival of peripheral macrophages after CSF-1R inhibition. We observed that the CSF-1R antagonist, PLX5622, exerted significant elimination of peripheral tissue macrophages, albeit with lower efficiency than in the CNS. Furthermore, we discovered that Mr BMT achieved replacement efficiencies comparable to tBMT in peripheral tissues such as the liver, kidney, lung, and spleen. Our findings suggested Mr BMT is a highly efficient strategy for allogeneic macrophage replacement in both the CNS and peripheral organs, and it could have potential clinical applications.

## Introduction

Tissue-resident macrophages, including microglia which are brain parenchyma resident macrophages, continuously monitor and clear pathogens throughout the lifespan. The new paradigm suggests that several tissue-resident macrophage populations are seeded during waves of embryonic haematopoiesis and self-maintain independently of BM contribution during adulthood[1–3]. Microglia and other tissue-resident macrophages play essential roles in the development, homeostasis, tissue repair, and disease defence both in the CNS and peripheral organs. Dysfunction or genetic mutations have been linked to a wide range of diseases, including neurodegeneration, autoimmune disorders, inflammation, and bone disease[4]. Allogeneic bone marrow and stem cell transplantation are effective options for the treatment of inherited diseases of the immune system, such as severe combined immunodeficiency (SCID)[5] and some autoimmune diseases, including systemic lupus erythematosus (SLE)[6], rheumatoid arthritis (RA)[7], type 1 diabetes (T1D)[8] and multiple sclerosis (MS)[9]. This approach involves replacing the recipient’s bone marrow and immune cells with healthy donor cells that can repopulate the immune system, leading to improved disease outcomes.

We previously developed a novel strategy for nonself microglia replacement by bone marrow transplantation at CNS-wide scale and named this procedure Mr BMT[10]. This involves first depleting the resident microglia through either genetic or pharmacological means, followed by transplantation of bone marrow cells that giving rise to new microglia. The existing microglia of recipient mice will be replaced by microglia-like cell derived from bone marrow cells[11, 12]. This strategy can be useful in cases where the resident microglia are dysfunctional and contribute to the development of neurodegenerative, such as Alzheimer’s disease (AD)[13], Parkinson disease (PD)[14] and Frontotemporal dementia (FTD)[15]. However, it is important to evaluate the potential effects of Mr BMT on peripheral tissue macrophages when considering this therapy for clinical translation, since the approach used for microglia ablation also have significant effects on the tissue macrophages and monocyte populations[16–18].

The current study observed highly effective tissue macrophage replacement in the liver, kidney, lung, and spleen after Mr BMT, indicating the potential of this therapy to have broad impacts on tissue macrophage populations throughout the peripheral organs. These encouraging results suggested that Mr BMT may be a promising cell therapy strategy for a wide range of diseases that affect both the central and peripheral immune systems, such as amyotrophic lateral sclerosis (ALS) and adrenoleukodystrophy (ALD).

## Materials and Methods

### Animals and PLX5622 administration

*Cx3cr1*-GFP (B6.129P2(Cg)-*Cx3cr1^tm1Litt^*/J, Strain #: 005582) mice were purchased from the Jackson Laboratory (Shanghai, China)[19]. C57BL/6J mice were purchased from Charles River (Shanghai, China). All mice were housed in the Animal Facility at the Department of Laboratory Animal Science at Fudan University under a 12 hours light/dark cycle with food and water given ad libitum. All animal experiments were conducted in accordance with the guidelines of the Institutional Animal Care and Use Committee of the Department of Laboratory Animal Science at Fudan University (202009001S and 2021JS-NZHY-002). All PLX5622 were formulated in AIN-76A chow at a concentration of 1200 mg/kg (Cat# D20010801, SYSE Bio, Changzhou, China). PLX5622 group mice were fed with PLX5622 chow for 14 days ad libitum. Control diet group mice were fed with AIN-76A chow ad libitum.

### Tissue preparation for the cryosection and immunohistochemistry

Mice were deeply anesthetized with isoflurane and were then subjected to transcranial perfusion with 50 ml of 0.01 M cold PBS or 0.9% NaCl. Next, all mice were perfused with 4% cold paraformaldehyde (PFA) for fixation. The brains and peripheral organs, including liver, lung, kidney and spleen were collected and postfixed in 4% PFA overnight (ON) at 4 °C or 6 hours in room temperature (RT). All tissues were dehydrated by 30% sucrose in 0.01 M PBS in 4 °C for 2-3 days and then embedded in optimal cutting temperature compound (OCT). All samples were stored at −80 °C before cryosection. All tissues were frozen in the freezing microtome for 15 minutes before cryosection. Tissues with regions of interest were sectioned by Leica CM1950 cryostat at 35 μm thickness.

### Immunohistochemistry and image acquisition

The sections of brain and peripheral organs were rinsed by 0.01 M PBS for 3 times. Next, brain and peripheral organ sections were blocked and permeabilized with 0.01 M PBS containing 0.03% Triton X-100 and 4% normal donkey serum (NDS) for 2 hours at RT. Next, the samples were incubated in primary antibody in 1% NDS in PBST at 4 °C ON. After rinsed by 0.03% PBST for 3 times, the samples were incubated fluorescent dyes conjugated secondary antibody in 1% NDS in PBST for 2 hours at RT, together with 1:500 4’, 6-diamidino-2-phenylindole (DAPI). Next, samples were rinsed by 0.03% PBST for 3 times and then mounted anti-fade mounting medium. The edge of the cover glass was sealed with nail enamel.

Primary antibody information and concentration: rabbit anti-IBA1 (1:500, Cat# 019-19741, Wako, Japan); goat anti-IBA1 (1:500, Cat# ab5076, Abcam, Cambridge, UK); goat anti-GFP (1:1000, Cat# ab6673, Abcam, Cambridge, UK); rabbit anti-GFP (1:1000, Cat# A11122, Thermo Fisher, Massachusetts, USA); rat anti-F4/80 (1:500, Biolegend, Cat# 123101, California, USA); mouse anti-CD11c (1:500, Cat# 117314, Biolegend, California, USA). Secondary antibody information: Donkey anti-rat 647 (1:1000, Cat# A48272, Invitrogen, California, USA); Donkey anti-rat 568 (1:1000, Cat# ab175475, Abcam, Cambridge, UK); Donkey anti-rabbit 488 (1:1000, Cat# A11008, Thermo Fisher, Massachusetts, USA); Donkey anti-goat 488 (1:1000, Cat# A11055, Thermo Fisher, Massachusetts, USA); Donkey anti-rabbit 568 (1:1000, Cat# A10042, Thermo Fisher, Massachusetts, USA); Donkey anti-rabbit 647 (1:1000, Cat# 711-605-152, Jackson, Pennsylvania, USA).

Confocal images were acquired by using an Olympus FV3000 confocal microscope with a solid-state laser. Lasers with wavelengths of 405 nm, 488 nm, 561 nm, and 640 nm were used to excite the fluorophores. Then, ×60 (oil), ×40 (oil), and ×20 objectives were utilized. Some whole-brain fluorescence images were acquired by an Olympus VS120 microscope equipped with a motorized stage. Then, ×10 objective was used. Z-stacked focal planes were acquired and maximally projected with Fiji. The brightness and contrast of the image were adjusted with Fiji if necessary.

### Bone marrow cells preparation

Bone marrow cells were collected from adult *Cx3cr1*^+/GFP^ mice. In brief, femora and tibia were collected and the muscles were removed. Next, the bones were placed in cold 0.01 M PBS and washed twice. After that, bone marrow cells were collected using 10 mL syringe with a 26-gague needle. Next, cells were washed by L15 medium containing 10% FBS and centrifuged at 300 g for 10 minutes. Cell pellets were then resuspended with L15 medium containing 10% FBS and passed through 40 μm cell strainers. After that, cell number was counted, and bone marrow cells were diluted in appropriate amount medium for downstream applications.

### Traditional bone marrow transplantation (tBMT)

The tBMT was carried out according to our previous description with some modifications[10]. Briefly, adult recipient C57BL/6J mice were fed with acid water (PH=2∼3) and neomycin (1.1 g/L) for 14 days till receiving 9 Gy X ray treatment. After irradiation, the recipient mice were administrated with GFP-labelled 1 × 10^7^ bone marrow cells from the donor through tail vein injection.

### Microglia replacement by bone marrow transplantation (Mr BMT)

The Mr BMT was carried out according to our previous description with some modifications[20]. In brief, adult recipient C57BL/6J mice were administrated PLX5622 together with acid water (PH=2∼3) and neomycin (1.1 g/L) for 2 weeks till receiving 9 Gy X ray treatment. At the same day, the recipient mice were intravenous administrated with GFP-labelled 1 × 10^7^ bone marrow cells from the donor through tail vein injection.

### Statistical analysis

Statistical analyses were conducted using Prism 9 (GraphPad). Each data point represented the average statistical result of three sections in different regions of brain, liver, kidney, lung and spleen. Results were determined independently in double-blind manner. Data were presented as mean ± standard deviation (SD). One-way analysis of variance (ANOVA) with Holm-Sidak’s multiple comparisons test (post hoc) was performed for multiple comparisons, while unpaired t test was performed to compare the differences between two groups, unless otherwise stated. Statistical significance was set as p < 0.05.

## Results

### CSF-1R inhibitor-induced elimination of tissue macrophages

The colony stimulating factor-1 (CSF-1) signalling pathway is known to regulate the development, proliferation, migration, function and survival of myeloid cells, including microglia and peripheral tissue macrophages[21]. The PI3K/Akt pathway has a central role in CSF-1R mediated macrophage survival. In macrophages, Akt can be activated directly through the CSF-1R pTyr721/PI3K pathway and indirectly by ceramide-1-phosphate (C1P) or the Gab2/PI3K pathway, which is counterbalanced by a CSF-1R pTyr559/LynSHIP-1 pathway[22]. Previous study showed that the inhibition of CSF-1R would result in the elimination of microglia in the adult brain and the establishment of a microglia-free niche[23]. Meanwhile, the survival of peripheral tissue-resident macrophages and monocyte are also threatened[18]. Additionally, interference with CSF-1R signalling through depleting *Csf1r* enhancer can lead to the lack of myeloid cells, including the microglia in brain and macrophages in skin, kidney, heart, and peritoneum. However, the other periphery tissue macrophages are unaffected[24].

The small molecule PLX5622 is a CSF-1R antagonist that has been tested in a range of animal models[23, 25, 26]. Here, adult mice were administered control and PLX5622 chow for 14 days to examine whether CSF-1R inhibition could generate a macrophage-free niche in peripheral tissues within this time window (Fig. 1A). IBA1 was used as a marker to determine the number of microglia in brain and macrophages in peripheral organs, as IBA1 is specifically expressed in most subpopulations of monocyte and macrophage lineage cells[27]. Consistent with previous studies, we observed that PLX5622 treatment resulted in the elimination of ∼99% of microglia and the creation of a microglia-free environment in the brain[23]. However, unlike with the observation in brain, 44.08% IBA1-positive macrophages were depleted in the liver, 50.58% in the lung, 87.123% in the kidney, 31.41% in the spleen after 14 days PLX5622 treatment (Fig. 1B and Fig. S1). These observations demonstrated that PLX5622 exerted a significant elimination of peripheral tissue macrophages, although its efficiency was lower than in the CNS.

**Fig. 1.**
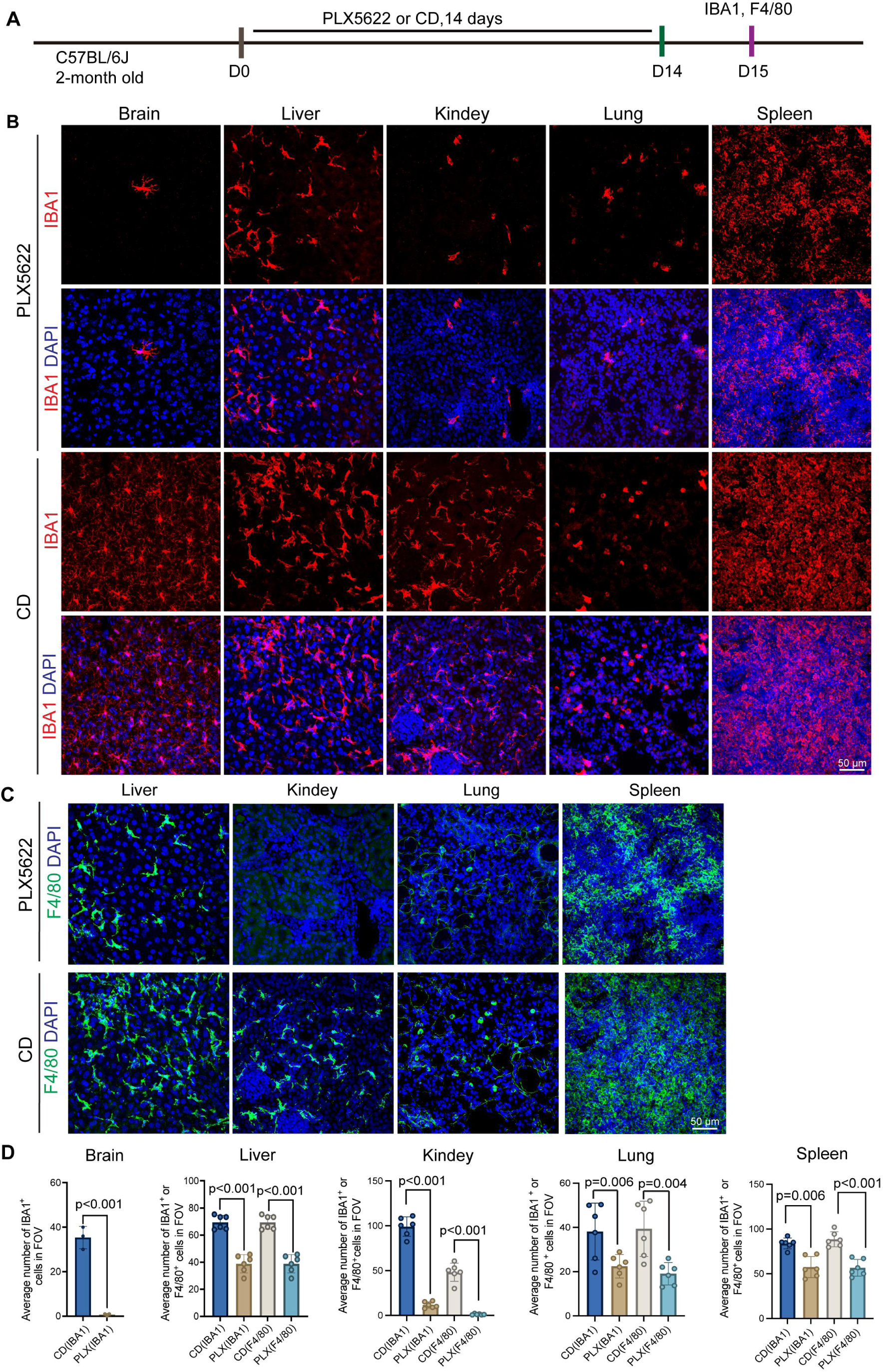
The CSF-1R antagonist, PLX5622, depleted microglia in the brain and resident macrophages in peripheral tissue. (A) Scheme of time points for microglial depletion and examination time points. (B) Representative images show the depletion efficiency of microglia (IBA1, red) in the brain or peripheral macrophages (IBA1, red) in the peripheral organs following the administration of CD or PLX5622. Scale bar, 50 μm. (C) Representative images show the depletion efficiency of tissue macrophages (F4/80, green) in peripheral organs following the administration of PLX5622. Scale bar, 50 μm. (D) Quantification the average number of IBA1^+^ cells or F4/80^+^ cells in the indicated organs. N = 3∼6 mice. Two-tailed unpaired test. Data are presented as mean ± SD. CD: control diet; FOV: Field of View; D: day.

Although F4/80 is widely used in the analysis of macrophages, it is also expressed in other immune cells, such as dendritic cells (DCs) in the spleen[28]. We evaluated the co-expression rate of F4/80 and IBA1 in various tissues. Our results showed that all F4/80^+^ cells expressed IBA1 in the liver, but 85.67% F4/80^+^ cells expressed IBA1 in the lung, and 50.17% in the kidney, 92.96% in the spleen (Fig. S2). Additionally, the elimination rate of F4/80-positive cells after PLX5622 treatment varied among tissues, with 44.08% in the liver, 51.62 % in the lung, 97.46% in the kidney, and 36% in the spleen (Fig. 1C and 1D). Interestingly, the absent of F4/80^+^ macrophages in kidney after CSF-1R antagonist treatment was consistent with the findings in *Csf1* enhancer deletion mice[29]. Noticeably, granulocyte-macrophage colony stimulating factor 2 (GM-CSF), also known as colony stimulating factor-2 (CSF-2) regulates lung macrophage survival together with CSF-1[30]. CSF-2/CSF2RB signalling pathway is also important for alveolar macrophage survival, so inhibiting CSF-1R alone will not kill all alveolar macrophages. As CSF-1/CSF-1R signal pathway is crucial for the proliferation and survival of patrolling monocytes[31, 32], the declined number of IBA1 and F4/80 cells in the lung after PLX5622 administration was also likely attributed to the elimination of monocytes.

### Mr BMT successfully replaced Kupffer cells with BM-derived macrophages in the liver

We previously discovered that microglia-free niche is a prerequisite condition for the successful engraftment of BM-derived cells[10]. We thus hypothesized that the elimination of tissue macrophages by utilization of CSF-1R inhibitor would improve the BM engraftment efficiency. To evaluate the replacement efficiency of macrophages in the liver, we used *Cx3cr1*^+/GFP^ mice as donor, as CX3CR1 is expressed in monocyte/macrophage lineage cells[19]. The recipient mice were fed with PLX5622-formulated chow for 14 days and then subjected to X-ray irradiation, followed by an intravenous injection of 1×10^7^ BMCs from *Cx3cr1*^+/GFP^ as we previously described[11]. After that, the recipient mice were fed with control diet for thirty days to remove the CSF-1R inhibition (Fig. 2A). We replaced 98.62% of brain resident microglia with BM-derived microglia-like cells (Fig. S3), indicating that Mr BMT exhibits an exceptionally high level of transplantation effectiveness within the brain. We firstly examined BM-derived macrophages ratio in the peripheral organs of the chimeric mouse, then the morphological changes were also included in the analysis.

**Fig. 2.**
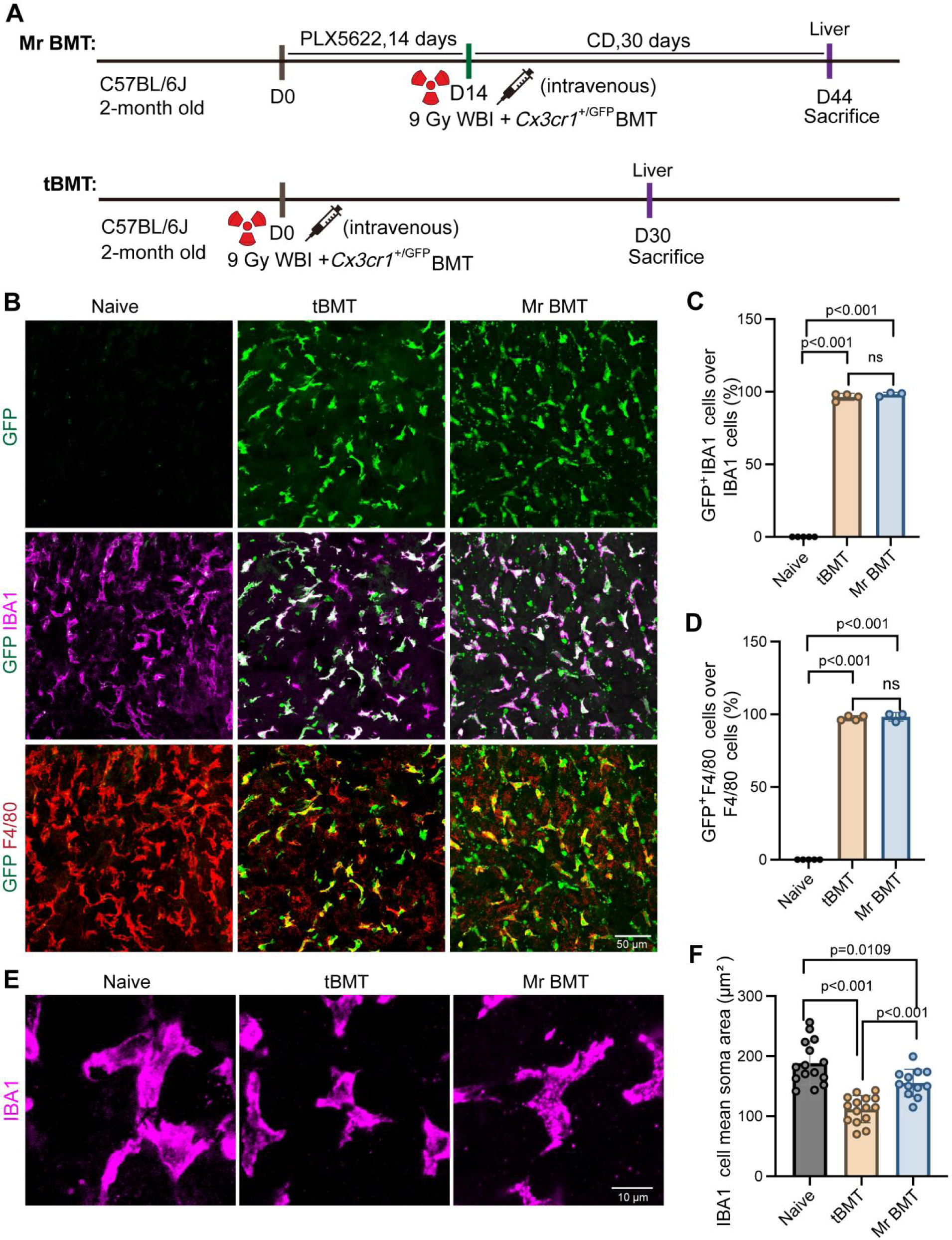
Mr BMT successfully replaced Kupffer cells with BM-derived macrophages in the liver. (A) Scheme of time points for Mr BMT and tBMT strategy. (B) Representative images show BM-derived cells (GFP, green), liver resident macrophages (IBA1, magenta), liver resident macrophages (F4/80, red) in the liver after tBMT or Mr BMT treatment. Scale bar, 50 μm. (C-D) Quantification the percentage of tBMT-derived cells and Mr BMT-derived cells in the liver. N = 3∼5 mice. One-way ANOVA with Holm-Sidak multiple-comparisons test (post hoc). Data are presented as mean ± SD. (E) Representative images show the morphology of BM-derived KCs (IBA1, magenta) in the liver. Scale bar, 10um. (F) Quantification of soma area of IBA1^+^ cells in the liver. N = 3∼5 mice. For each mouse, take three non-consecutive sections, and each dot represents one cell. Each dot represents one cell. One-way ANOVA with Holm-Sidak multiple-comparisons test (post hoc). Data are presented as mean ± SD. CD: control diet; WBI: whole brain irradiation; D: day.

In the adult mouse liver, Kupffer cells (KCs) are originated from yolk sac (YS) and fetal liver monocyte-derived macrophages and replenished independently by monocyte-derived precursors in the steady state[1]. Both cell populations share common macrophage functions, including phagocytosis, pathogen defence, native and adaptive immune regulation. Here, we observed that more than 98.22% of primary KCs were replaced by BM-derived GFP^+^IBA1^+^ macrophages at thirty days post Mr BMT. There was no significant difference in transplantation efficiency between tBMT and Mr BMT (Fig. 2B-2D). Compared with primary KCs, the body size of the BM-derived KCs was smaller, implying different cell originations of KCs are linked to the discrepancies in cell morphology and function like immune response and tissue repairment ability (Fig. 2E and 2F).

### Mr BMT strategy showed high replacement efficiency of kidney tissue macrophages

We further tested the macrophage replacement efficiency in the kidney after tBMT and Mr BMT. In the renal, the majority tissue macrophages are derived from YS and fetal liver macrophages, and the YS derived macrophages progressively expand in the mouse kidney with age[33]. These populations are maintained in situ by self-renewal, largely independent of adult haematopoiesis across lifespan[34, 35]. Immunofluorescent (IF) staining of GFP in kidney sections from recipient C57BL/6J mice revealed GFP^+^ BM-derived macrophages spread out the cortex and medulla after tBMT and Mr BMT (Fig. 3A). About 87.22% of tissue macrophages were reconstituted by GFP^+^IBA1^+^ BM-derived macrophages thirty days after tBMT (Fig. 3B and 3C). After recovery from Mr BMT, more than 87.50% of primary kidney macrophages were successfully colonized by GFP^+^IBA1^+^ BM-derived macrophages (Fig. 3B-3C), suggesting Mr BMT achieved a similar replacement efficiency to tBMT. Both in tBMT and Mr BMT group, the replacement rate of GFP^+^F4/80^+^ / F4/80^+^ was similar to GFP^+^IBA1^+^ / IBA1^+^ (Fig. 3B-3D). Previous study demonstrated the endogenic macrophages presented a distinct morphology and function compared to BM-derived macrophages[20, 36, 37]. Therefore, we examined the morphology of the newly formed macrophages in the kidney after tBMT and Mr BMT. We observed that the surface area was reduced significantly in BM-derived macrophages, implying BM-macrophages did not fully acquire a resident macrophage phenotype (Fig. 3E and 3F).

**Fig. 3.**
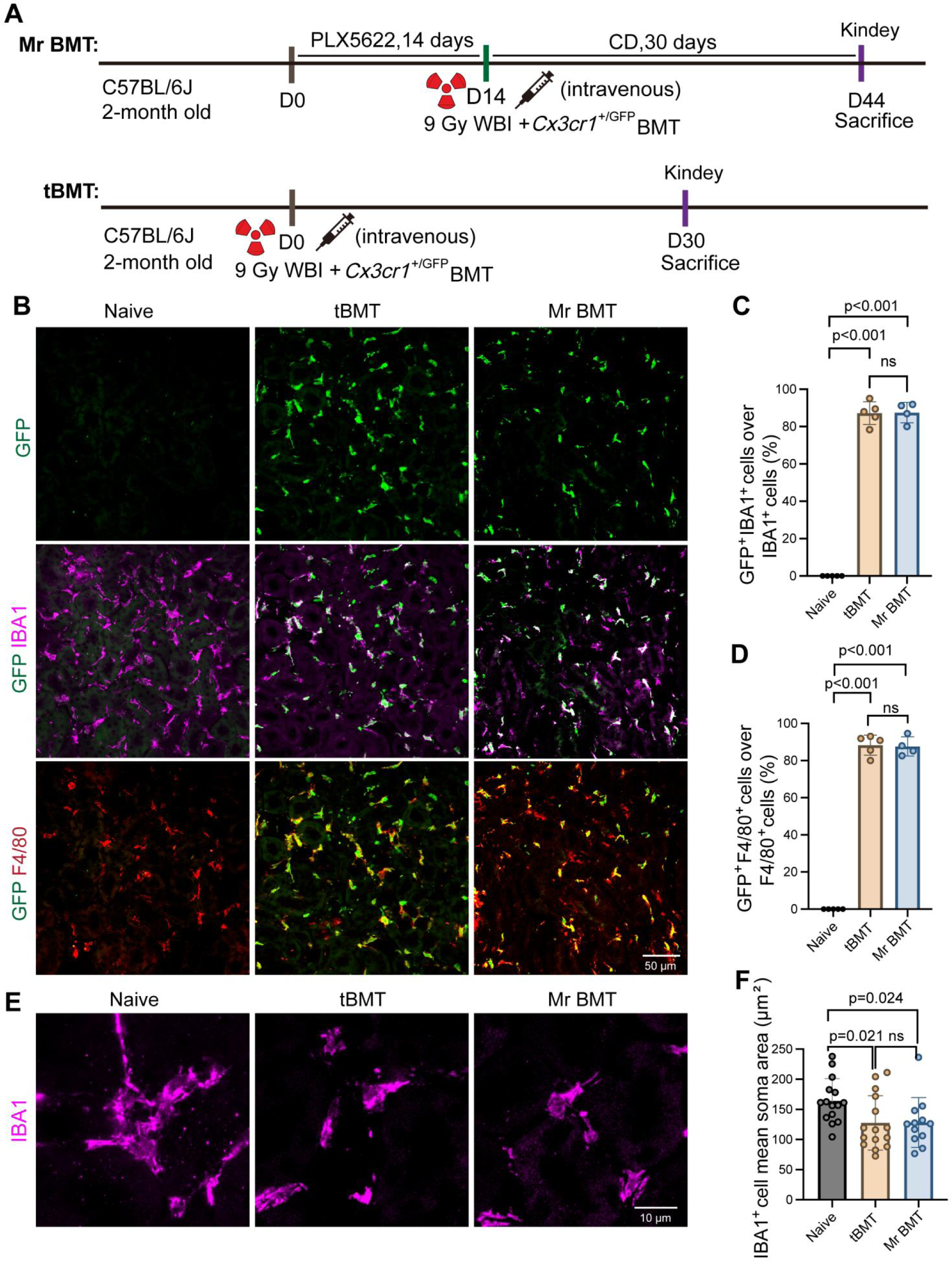
Evaluate the efficiency of the Mr BMT strategy in replacing macrophages in kidney. (A) Scheme of time points for Mr BMT and tBMT strategy. (B) Representative images show BM-derived cells (GFP, green), kidney resident macrophages (IBA1, magenta), kidney resident macrophages (F4/80, red) in the kideney after tBMT or Mr BMT treatment. Scale bar, 50 μm. (C-D) Quantification the percentage of tBMT-derived cells or Mr BMT-derived cells in the kidney. N = 4∼5 mice. One-way ANOVA with Holm-Sidak multiple-comparisons test (post hoc). Data are presented as mean ± SD. (E) Representative images showing the morphology of BM-derived kidney macrophages (IBA1, magenta) in the kidney. Scale bar, 10 μm. (F) Quantification of soma area of IBA1^+^ cells in the kidney. N = 4∼5 mice. For each mouse, take three non-consecutive sections, and each dot represents one cell. Each dot represents one cell. One-way ANOVA with Holm-Sidak multiple-comparisons test (post hoc). Data are presented as mean ± SD. CD: control diet; WBI: whole brain irradiation; D: day.

### Mr BMT achieved high engraftment efficiency of macrophages in the lung and spleen

Next, we examined the profile of BM-originated macrophages in the lung either after tBMT or Mr BMT (Fig. 4A). After transplantation, a great number of GFP^+^ BM-derived cells proliferated in the recipient mouse lung, and subpopulations of them were labelled by macrophage/monocyte lineage marker IBA1 and F4/80. After recovery from tBMT, about 80.70% endogenous macrophages/monocytes cells were replaced by GFP^+^ F4/80^+^ BM cells. In Mr BMT group, there were 85.17% IBA1^+^ and 76.53% F4/80^+^ cells in the recipient mouse lung were detected as BM-derived GFP^+^ cells after Mr BMT (Fig. 4B-4D), suggesting comparable replacement efficiency to tBMT. Previous study has demonstrated that CX3CR1^+^ BMCs could give rise to DCs and supply tissue DCs in lung[38, 39]. We found 2.81% of GFP^+^ cells that were IBA1^-^ and F4/80^-^, which were likely to differentiate into pulmonary DCs (Fig. 4E and 4F). Lung resident macrophage contained alveolar macrophages (AMs) and interstitial macrophages (IMs)[40]. AMs have high expression of CD11c and low expression of CX3CR1[41, 42], which is totally different from the GFP^+^ cells after tBMT or Mr BMT. Thus, we hypothesized that the replacement cells more likely to convert to IMs after thirty days recovery.

**Fig. 4.**
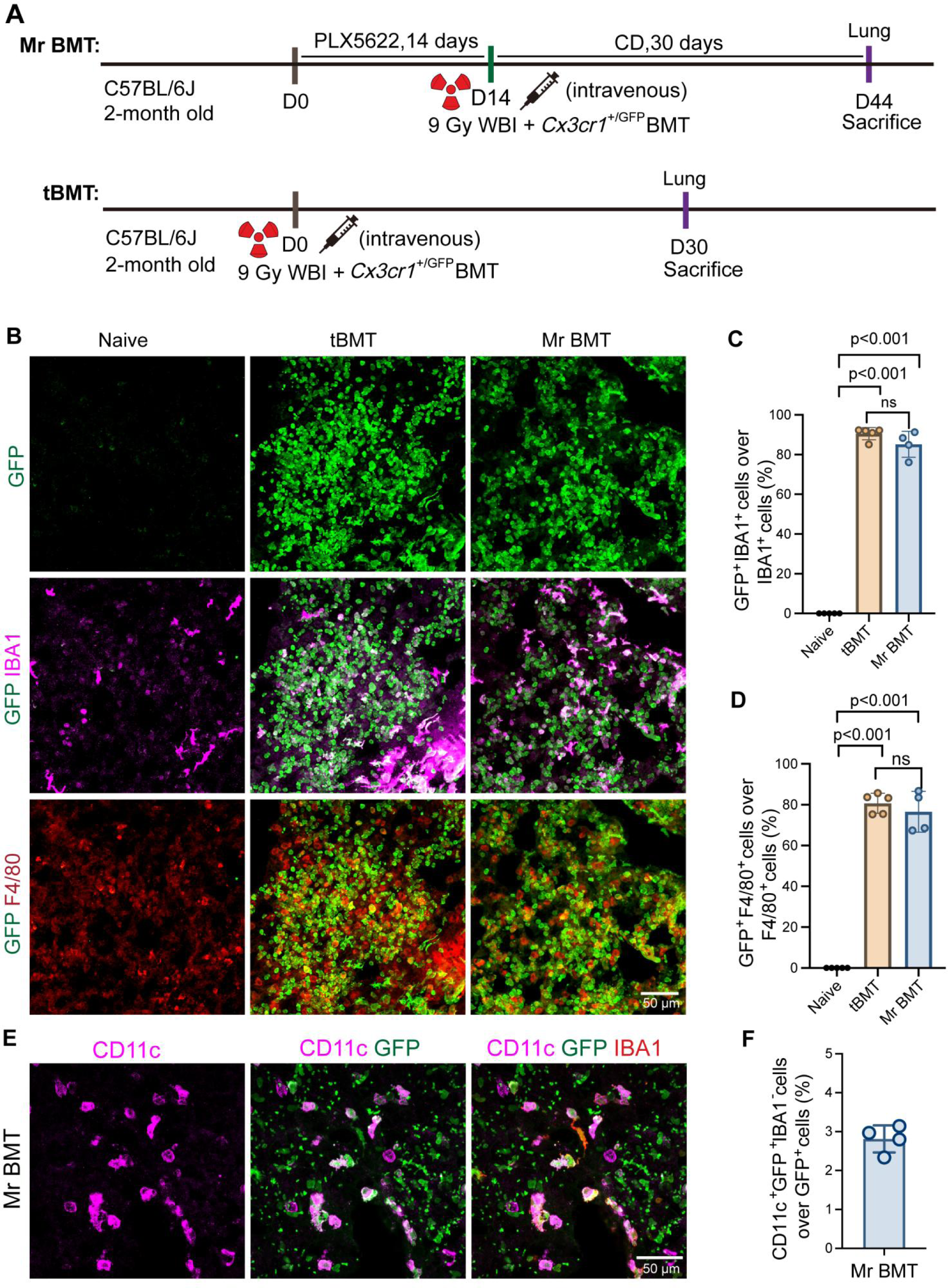
Evaluate the replacement efficiency of lung tissue-resident macrophages under Mr BMT treatmet. (A) Scheme of time points for Mr BMT and tBMT strategy. (B) Representative images show BM-derived cells (GFP, green), lung resident macrophages (IBA1, magenta), lung resident macrophages (F4/80, red) in the lung after tBMT or Mr BMT treatment. Scale bar, 50 μm. (C-D) Quantification the percentage of tBMT-derived cells or Mr BMT-derived cells in the lung. N = 4∼5 mice. One-way ANOVA with Holm-Sidak multiple-comparisons test (post hoc). Data are presented as mean ± SD. (E) Representative images show myeloid cells (CD11c, magenta), BM-derived cells (GFP, green) and lung tissue-resident macrophages (IBA1, red) in the lung after Mr BMT. Scale bar, 50 μm. (F) Quantification the ratio of CD11c^+^ GFP^+^ IBA1^-^ cells / GFP^+^ cells in the lung after Mr BMT. N = 4 mice. Data are presented as mean ± SD. CD: control diet; WBI: whole brain irradiation; D: day.

Lastly, we investigated the replacement efficiency of macrophages/monocytes in the spleen after received *Cx3cr1*^+/GFP^ transplantation (Fig. 5A). Regardless of the approaches used for BMT, the staining results revealed that more than 50% of tissue-resident macrophages were replaced by GFP^+^ BM-derived cells (Fig. 5B-5D). Compared with other peripheral organs, the replacement rate in the spleen was lower, which may be attributed to the different macrophage turnover rate in different organs. However, similar with the observations in other peripheral organs, BM-derived macrophages in the spleen possessed a smaller cell body and less branches, differing from the endogenous embryonic originated macrophages. Additionally, there are 6.96 % GFP^+^ cells that were IBA1^-^ and F4/80^-^, which were likely to contribute to DCs in the spleen (Fig. 5E and 5F).

**Fig. 5.**
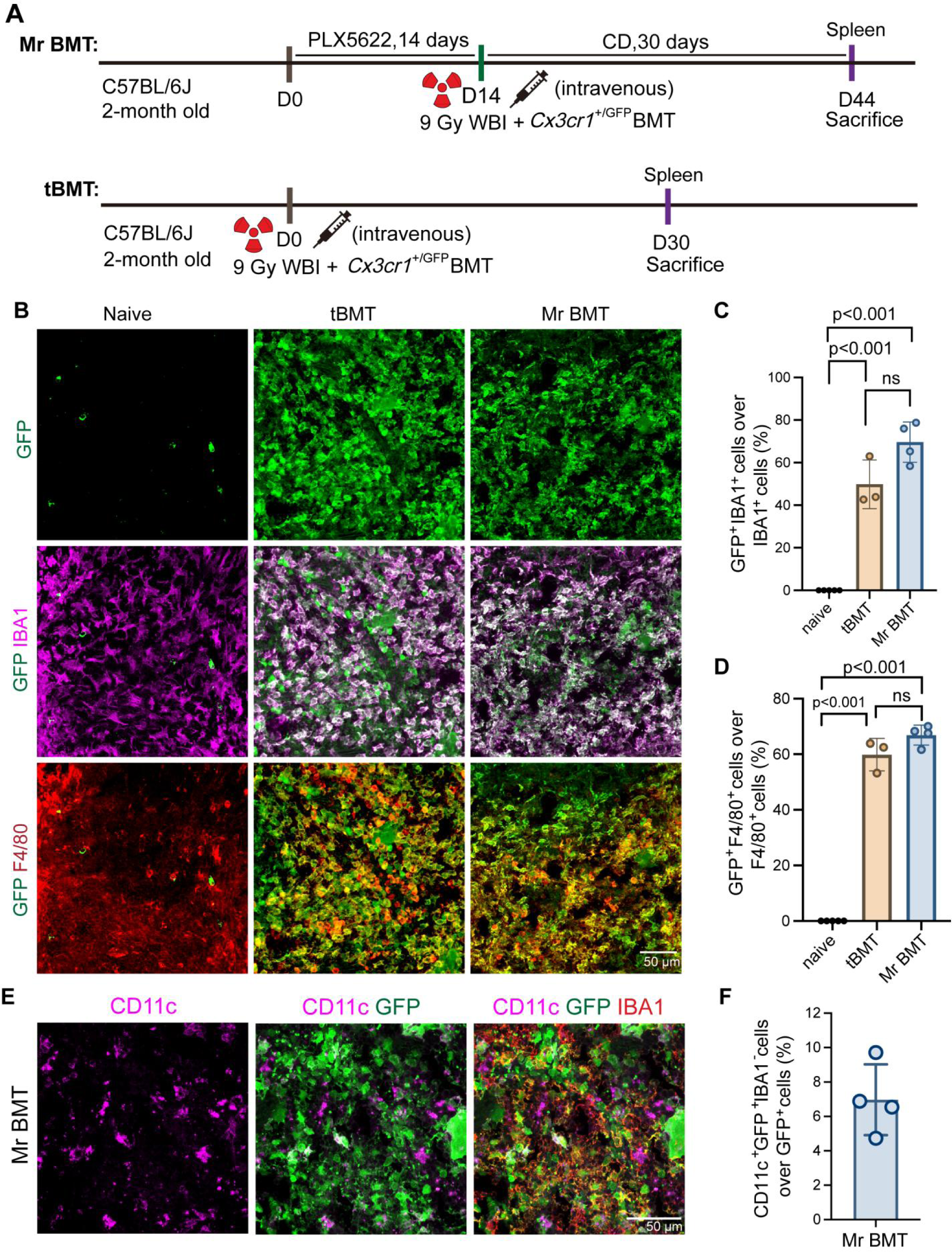
Replacement efficiency of spleen tissue-resident macrophages under Mr BMT treatmet. (A) Scheme of time points for Mr BMT and tBMT strategy. (B) Representative images show BM-derived cells (GFP, green), spleen resident macrophages (IBA1, magenta), spleen resident macrophages (F4/80, red) in the liver after tBMT or Mr BMT treatment. Scale bar, 50 μm. (C-D) Quantification the percentage of tBMT-derived cell or Mr BMT cell percentage in the spleen. N = 3∼5 mice. One-way ANOVA with Holm-Sidak multiple-comparisons test (post hoc). Data are presented as mean ± SD. (E) Representative images show myeloid cells (CD11c, magenta), BM-derived cells (GFP, green) and spleen tissue-resident macrophages (IBA1, red) in the spleen after Mr BMT. Scale bar, 50 μm. (F) Quantification the ratio of CD11c^+^ GFP^+^ IBA1^-^ cells / GFP^+^ cells in the spleen after Mr BMT. N = 4 mice. Data are presented as mean ± SD. CD: control diet; WBI: whole brain irradiation; D: day.

## Discussion

The Mr BMT approaches a highly efficient strategy for the replenishment of endogenous microglia with BM-derived microglia-like cells at the CNS-wide scale (over 92%) in adult mouse. In contrast, BMCs do not extensively proliferate and engraft the recipient brain (less than 5∼20%) by using tBMT method. In current study, we further investigated tissue-resident macrophages replacement efficiency in the peripheral organs by using the Mr BMT approach. Our results demonstrated that BM-derived macrophages efficiently reconstituted tissue macrophages in the liver, kidney, lung and spleen. Although the replacement efficiency had a subtle difference in different organs, which may be attributed to the different turnover rate of tissue-resident macrophages. Thus, Mr BMT achieved a similarly high replacement efficiency comparable to tBMT, but owned higher efficiently in the CNS. These findings suggested that the Mr BMT approach could be a powerful tool for allogenic macrophages replacement in both CNS and peripheral organs. It means that Mr BMT may be a promising cell therapy strategy for a wide range of diseases that affect both central and peripheral myeloid cells, such as ALS and ALD. Despite these encouraging findings, it is important to fully understand the safety of Mr BMT, including its potential long-term effects on tissue macrophage populations and any risks associated with the irradiation used for myelosuppression. Nonetheless, with continued research and development, the Mr BMT strategy may ultimately offer a powerful tool for the treatment of a range of diseases that affect the immune system.

### Ethics approval and consent to participate

All animal experiments were conducted in accordance with the guidelines of the Institutional Animal Care and Use Committee of the Department of Laboratory Animal Science at Fudan University (permit number: 202110005S).

## Declaration of Competing Interest

The authors have no commercial, proprietary or financial interest in the products or companies described in this article.

## Author contributions

Y.R. supervised and conceptualized this study. Y.R., and B.Y. wrote the manuscript. B.Y. and Y.H. performed most experiments. B.Y. analysed the staining results, J.D. and Y.H. provided necessary study support. All authors discussed the results and commented on this manuscript. And at last, we appreciate Open AI’s assistance in language.

## Acknowledgements

The authors thank Yanxia Rao and Bo Peng (Fudan University) for their sponsorship and support of this experiment. Fang Lei and Jiachen Zhu (Fudan University) for the excellent laboratory management. In addition, the authors express their gratitude and respect to all animals sacrificed in this study.

## Data Availability Statement

The raw data supporting the conclusion of this article will be made available by the authors, without undue reservation.

---

**Fig. S1.**
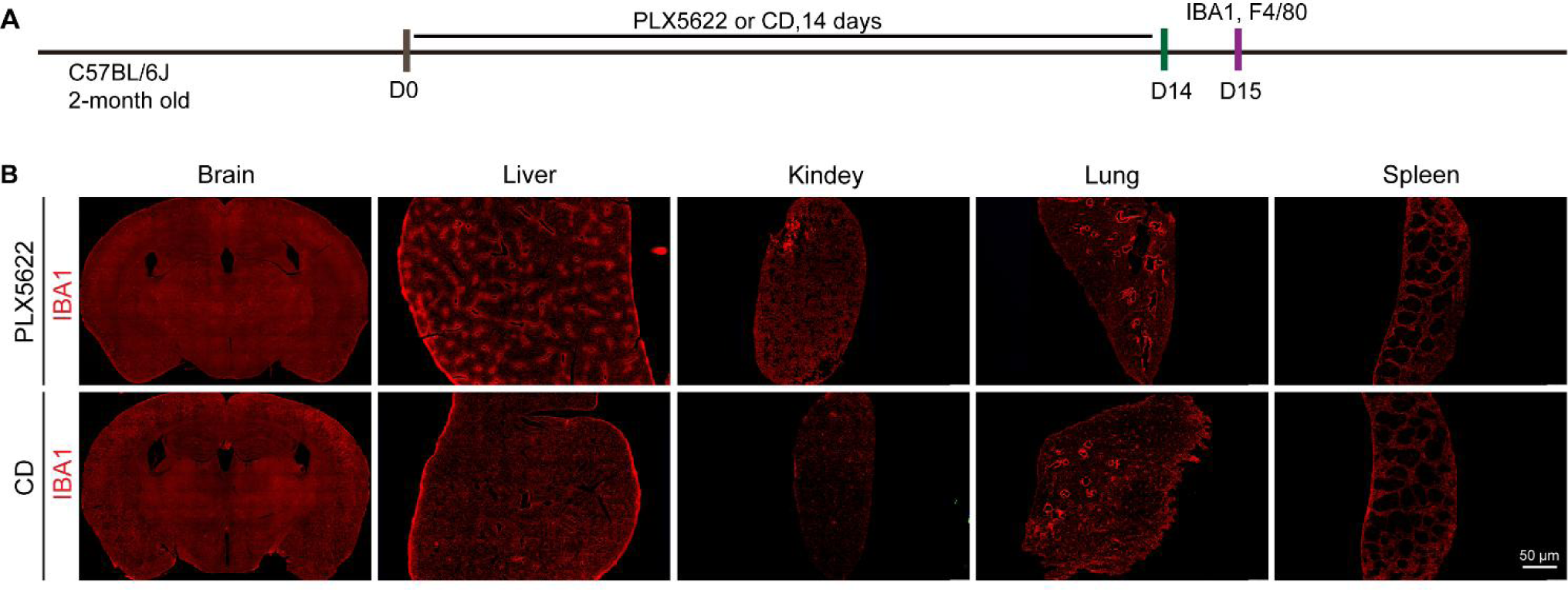
PLX5622 depleted microglia in CNS and resident macrophages in peripheral organs. (A) Scheme of time points for microglial depletion and examination time points. (B) Representative images show the depletion efficiency of microglia (IBA1, red) in the whole brain or macrophages (IBA1, red) in the peripheral organs following the administration of CD or PLX5622. Scale bar, 50 μm. CD: control diet; D: day.

**Fig. S2.**
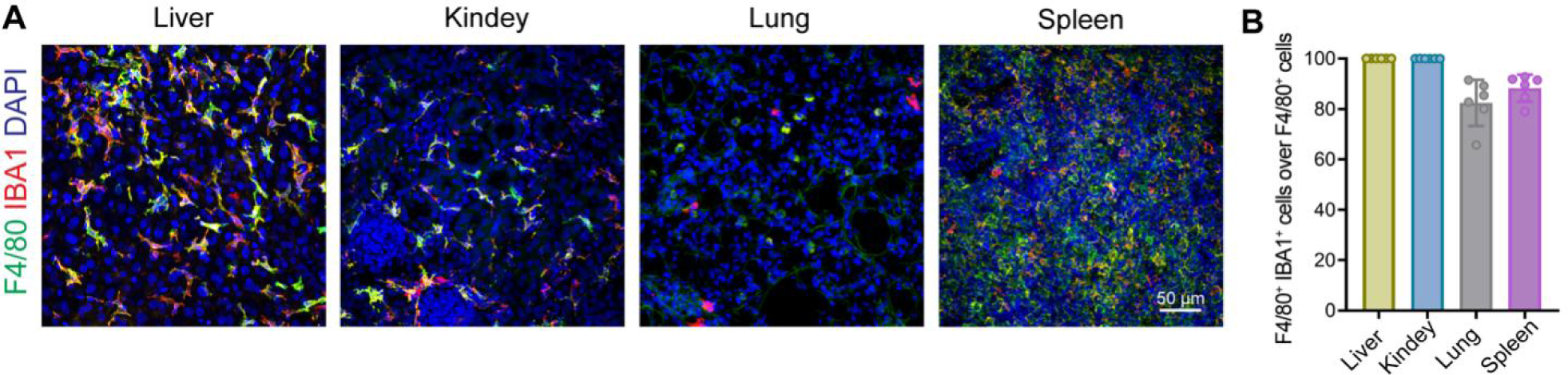
The co-lable rate of F4/80 and IBA1 in peripheral tissues. (A) Representative images show the co-labeling of F4/80 and IBA1 in the peripheral organs of naïve group. (B) Quantification the percentage of F4/80^+^ IBA1^+^ cells over IBA1^+^ cells in different peripheral tissues. N = 6 mice. Data are presented as mean ± SD. Scale bar, 50 μm.

**Fig. S3.**
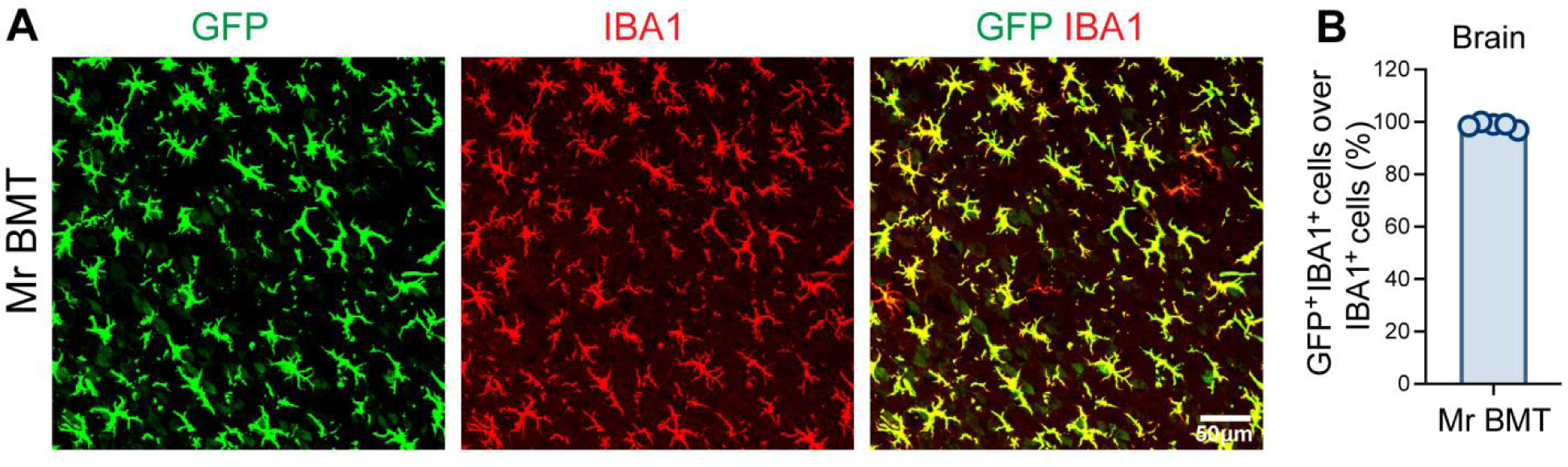
The replacement efficiency of Mr BMT in the brain was higher than in peripheral. (A) Representative images show the engrafted BM-derived microglia-like cells (GFP, green) in the brain after Mr BMT. Scale bar, 50 μm. (B) Quantification the percentage of Mr BMT-derived cells in the brain. N =5 mice. Data are presented as mean ± SD.

## Notes

### Competing Interest Statement

The authors have declared no competing interest.

### Summary of Updates

Change total number of pages: 34; word counts for abstract: 187; for the text: 6019

